# Reconciling the effects of PMS2 in different repeat expansion disease models supports a common expansion mechanism

**DOI:** 10.1101/2024.08.13.607839

**Authors:** Diego Antonio Jimenez, Carson J. Miller, Alexandra Walker, Kusala Anupindi, Bruce E. Hayward, Hernan A. Lorenzi, Karen Usdin, Xiaonan Zhao

## Abstract

Expansion of a disease-specific tandem repeat is responsible for >45 Repeat Expansion Diseases (REDs). The expansion mutation in each of these diseases has different pathological consequences and most are currently incurable. If the underlying mechanism of mutation is shared, a strategy that slows repeat expansion in one RED may be applicable to multiple REDs. However, the fact that PMS2, a component of the MutLα mismatch repair complex, promotes expansion in some models and protects against it in others suggests that the expansion mechanisms may differ. We show here using mouse models of two REDs caused by different repeats that PMS2 has similar effects in both models, with the loss of PMS2 resulting in an increase in expansions in some tissues and a loss of expansion in others. This is consistent with a protective effect of PMS2 in the first case and a role in promoting expansion in the second. Furthermore, we show in mouse embryonic stem cells that lower levels of PMS2 promote expansion while higher levels protect against it, with the ability to promote expansion depending on the PMS2 nuclease domain. Our findings lend support to the hypothesis that REDs share a common expansion mechanism and provide insights into the processes involved.

**Significance statement:** Collectively the Repeat Expansion Diseases (REDs) represent a significant health burden. Since the consequences of the expansion mutation differ across diseases, therapeutic approaches that block the underlying mutation are appealing, particularly if the mechanism is shared. However, the conflicting effects of PMS2 loss in different RED models challenges this idea. Here we show using two different RED models that these disparate effects can be reconciled into a single model of repeat expansion, thus increasing confidence that the REDs do all share a common mutational mechanism.

## Introduction

Repeat expansion, the increase in the number of repeats in a short tandem repeat (STR) in a disease-specific gene, is the cause of the Repeat Expansion Diseases (REDs). REDs are a group of >45 life-limiting neurological or neurodevelopmental disorders (1) with a collective allele frequency of 1 in 283 individuals (2). Converging evidence from studies of genetic modifiers in patients with different REDs implicates components of the Mismatch Repair (MMR) pathway as modifiers of both repeat expansion and disease severity (3-7). This has raised the possibility that targeting some of these factors may be useful therapeutically (8), an appealing idea since these diseases currently have no effective treatment or cure. Furthermore, if indeed these diseases share a common mechanism, a single treatment may be useful for multiple diseases in this group. Some of the same factors identified as modifiers of somatic expansion in human studies have been implicated in expansion in different cell and mouse models of these disorders. For example, MSH3, the MSH2-binding partner in MutSβ, one of the two lesion recognition complexes involved in mismatch repair (MMR), is required for expansion in multiple disease models (9-12). The same is true for MLH3 (13-18), the binding partner of MLH1 in the MutLγ complex involved in lesion processing in MMR (19). Furthermore, we and others have shown that the MLH3 nuclease domain is required for MLH3’s role in expansion (13, 18, 20). In MMR, MutSβ binds small insertions/deletions (IDLs) (21) and recruits either MutLγ or more commonly, Mutlα, a heterodimer of MLH1 and PMS2. MLH3 and PMS2 are endonucleases that introduce nicks in the substrate that ultimately leads to proper DNA repair (22). A third MLH1-binding partner, PMS1, has also been implicated in the generation of expansions in multiple disease models (14, 15, 23-25). However, PMS1 lacks the endonuclease domain seen in PMS2 and MLH3 and the MLH1-PMS1 heterodimer, Mutljβ, is not generally considered to be involved in MMR. However, recent work has shown that *in vitro* MutLβ is able to promote MutLγ-mediated expansions acting in part via stimulation of MutLγ cleavage (26).

While MLH3 and PMS1 both contribute to the generation of expansions in all model systems studied to date, PMS2 actually protects against expansions in some model systems while promoting expansion in others. For example, PMS2 protects against expansion in the brain (24, 25) and liver (27) of mouse models of Huntington’s disease (HD), a CAG-repeat expansion disorder, as well as in the brain of a mouse model of Friedreich’s ataxia (FRDA) (28), a GAA repeat expansion disorder (29). However, in a mouse model of Myotonic Dystrophy Type 1 (DM1), a CTG-repeat expansion disorder (30), loss of PMS2 results in the loss of ∼50% of expansions in many organs (31). In addition, we have shown that PMS2 is required for expansion in embryonic stem cells (ESCs) from a mouse model of the Fragile X-related disorders (FXDs) (23), disorders caused by a CGG-repeat expansion in the Fragile X Messenger Ribonucleoprotein 1 *(FMR1)* gene (32). This requirement for PMS2 was also seen in induced pluripotent stem cells (iPSCs) derived from a patient with Glutaminase Deficiency (GLSD) (14), another CAG-repeat expansion disorder (33). These differences raise the possibility that the expansion process differs in different REDs.

To address this possibility in a systematic way, we compared the effects of PMS2 loss in different organs of two different RED mouse models: a mouse model of the FXDs and a mouse model of **HD**. We also studied the effect of varying the expression of either normal PMS2 or PMS2 carrying a point mutation in its nuclease domain. Our findings have interesting implications for the expansion mechanism in different REDs.

## Results

### PMS2 plays a dual role in somatic expansion in an FXD mouse model

To examine the role of PMS2 in repeat expansion in an FXD mouse model, we crossed FXD mice to mice with a null mutation in *Pms2*. We then examined the expansion profiles in wildtype (WT), heterozygous and nullizygous animals matched for age and repeat number. Western blotting of proteins extracted from different tissues shows that heterozygosity results in significant loss of PMS2 protein in different tissues (Fig. S1). As can be seen in Fig. 1 and Fig. S2, this is associated with a small but significant increase in expansion in some tissues. In *Pms2* null mice, a further increase in expansion was seen in some tissues, including the striatum and cortex. However, in other organs, including colon and testes, fewer expansions were seen in *Pms2* null mice than were seen in heterozygous mice. Thus, in the very same animals, loss of PMS2 can both increase and decrease expansion depending on the tissue and the amount of PMS2 expressed.

**Figure 1.**
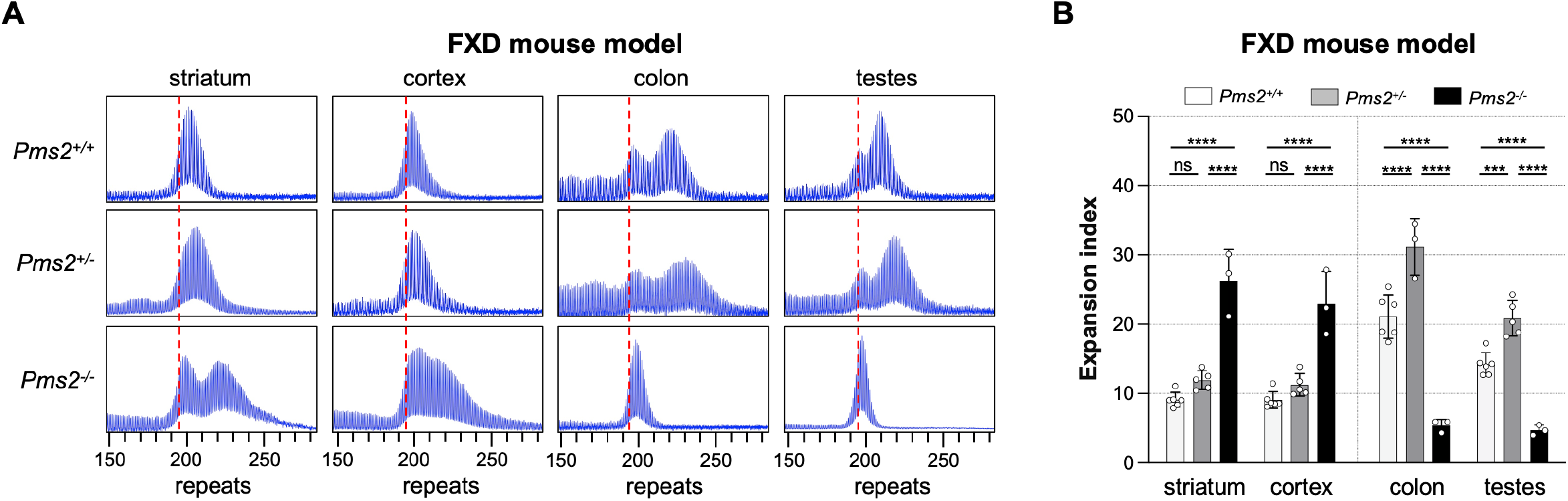
The effect of *Pms2* loss on repeat expansion in different tissues from an FXD mouse models. ((A) Representative repeat PCR profiles from striatum, cortex, colon and testes of 4-month-old *Pms2*^+/+^, *Pms2*^+/-^ and *Pms2*^-/-^ FXD male mice with 196 repeats. The dashed lines represent the sizes of the original inherited alleles as ascertained from the tail DNA taken at 3 weeks. (B) Comparison of the expansion index (EI) in the indicated organs of 4-month-old *Pms2*^+/+^, *Pms2*^+/-^ and *Pms2*^-/-^ FXD mice with an average of 194 repeats in the original allele. The colon data represent the average of 6 *Pms2*^+/+^, 3 *Pms2*^+/-^ and 3 *Pms2*^-/-^ mice with 185-210 repeats. The data from other organs represents the average of 6 *Pms2*^+/+^, 5 *Pms2*^+/-^ and 3 *Pms2*^-/-^ mice in the same repeat range. The error bars indicate the standard deviations of the mean. Each dot represents one animal. In each organ, the EIs for different genotypes were compared using a two-way ANOVA with correction for multiple testing as described in the Materials and Methods. The full tissue sets and the adjusted *p*-values are listed in the supplement Figure S2.***, *p* < 0.001; ****, *p* < 0.0001; ns: not significant.

### PMS2 plays a similar dual role in somatic expansion in an HD mouse model

To assess the role of PMS2 in repeat expansion in an HD mouse model, we crossed HD mice to the same *Pms2* null mice and again assessed repeat instability in animals matched for age and repeat number. As in the FXD mouse model, heterozygosity for *Pms2* results in an increase in expansions in many of the same tissues (Fig. 2 and Fig. S3). Furthermore, as in the FXD mice, in tissues such as striatum and cortex loss of all PMS2 resulted in a further increase in expansions, while in many other organs loss of all PMS2 resulted in a loss or decrease in expansions rather than an increase.

**Figure 2.**
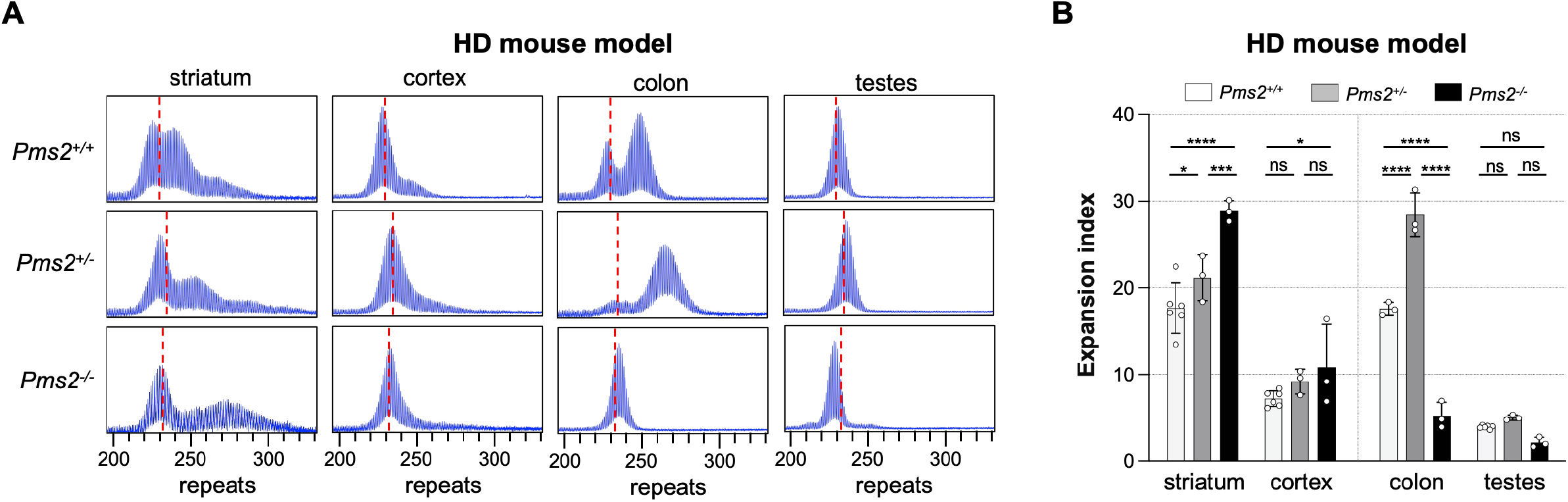
The effect of *Pms2* loss on repeat expansion in different tissues from an HD mouse model. (A) Representative repeat PCR profiles from striatum, cortex, colon and testes of 4-month-old *Pms2*^+/+^, *Pms2*^+/-^ and *Pms2*^-/-^ HD male mice with ∼230 repeats. The dashed lines represent the sizes of the original inherited alleles as ascertained from the tail DNA taken at 3 weeks. (B) Comparison of the expansion index (EI) in the indicated organs of 4-month-old *Pms2*^+/+^, *Pms2*^+/-^ and *Pms2*^-/-^ HD mice with an average of 234 repeats in the original allele. The colon data represent the average of 3 *Pms2*^+/+^, 3 *Pms2*^+/-^ and 3 *Pms2*^-/-^ mice with 226-239 repeats. The data from other organs represents the average of 6 *Pms2*^+/+^, 3 *Pms2*^+/-^ and 3 *Pms2*^-/-^ mice in the same repeat range. The error bars indicate the standard deviations of the mean. Each dot represents one animal. In each organ, the EIs for different genotypes were compared using a two-way ANOVA with correction for multiple testing as described in the Materials and Methods. The full tissue sets and the adjusted *p*-values are listed in the supplement Figure S2. *, *p* < 0.05; ***, *p* < 0.001; ****, *p* < 0.0001; ns: not significant.

### The differential effects of PMS2 on expansion are dependent on the PMS2 nuclease domain

To better understand the role of PMS2 in modulating expansions of the HD and FXD repeats we generated double knock-in (dKI) mESCs carrying∼180 FXD repeats and ∼215 HD repeats. These lines showed similar rates of expansion as single knock-in cell lines (Fig. 3A and D). We then used CRISPR-Cas9 to knock out *Pms2*. No expansion of either repeat was seen in the resultant lines (Fig. 3B, C, E, F). Thus, PMS2 is required for expansions of both repeats in this model system, consistent with our previous demonstration for the FXD repeat in mESCs (23) and the CAG repeat responsible for GLSD in patient iPSCs (7). As we had previously observed with other cell models (14, 23), complete loss of PMS2 resulted in small decreases in repeat number for both repeats. This is consistent with the idea that MLH3 levels in this cell type are too low to support expansion without a contribution from PMS2 and that loss of expansion then favors some sort of contraction.

**Figure 3.**
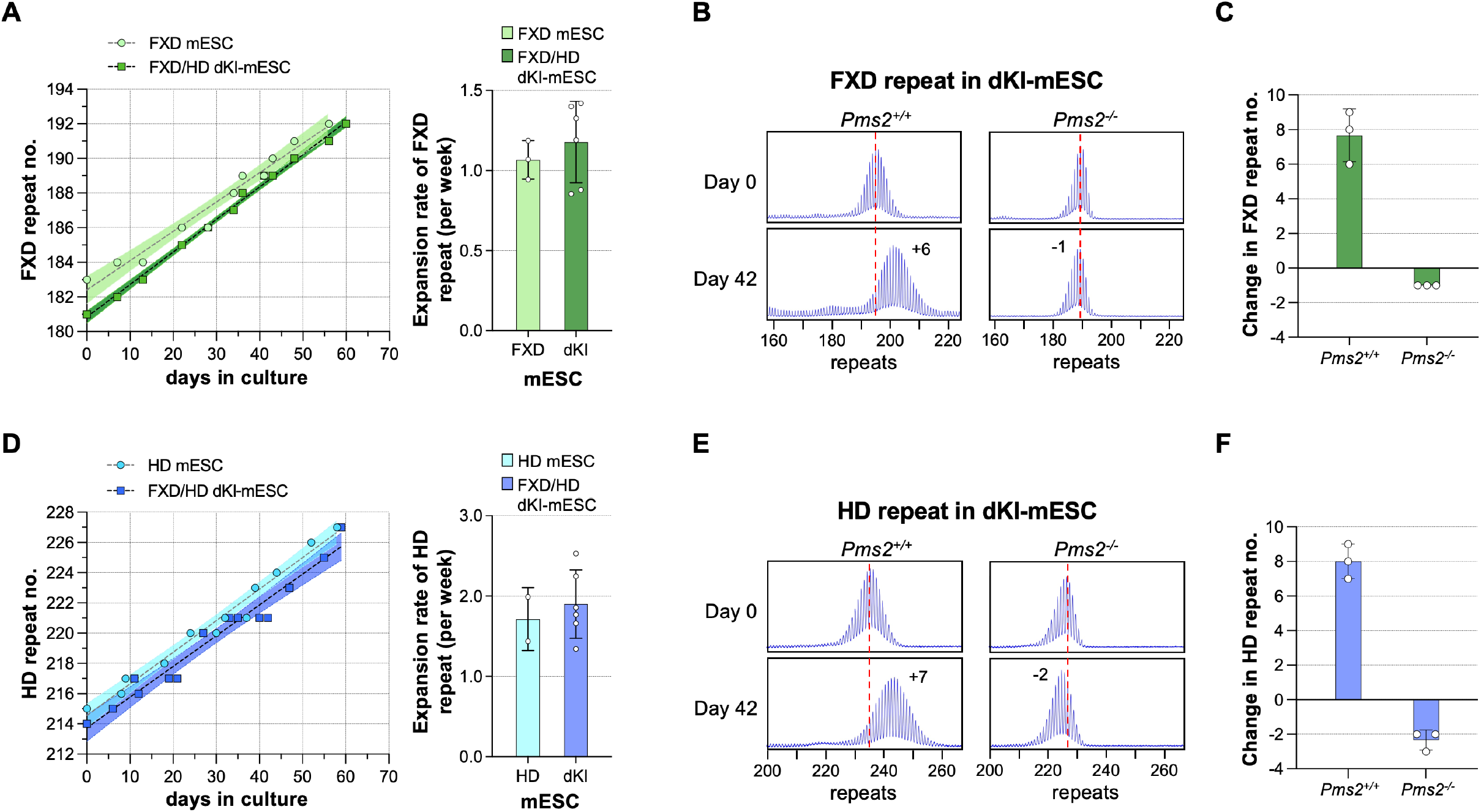
Effect of *Pms2* loss on expansion of FXD and HD repeats in a double knock-in mESC model. Representative graphs showing the change in repeat number over time for the FXD (A) and HD (D) repeats in either single or FXD/HD double knock-in (dKI) mESCs. The shaded area indicates the 95% confidence interval. Comparison of expansion rates of FXD and HD repeats in single and dKI mESCs are shown on the left. The expansion rate was calculated by linear regression and compared using a two-tailed unpaired t-tests as described in the Materials and Methods. The data represent the average of three FXD mESC lines with 183-186 repeats, two HD mESC lines with 212-215 repeats, and six FXD/HD dKI mESC lines with 172-194 FXD repeats (average182 repeats) and 206-233 HD repeats (average 220 repeats). The error bars indicate the standard deviations of the mean. Each dot represents one cell line. There is no significant di\erence between the rate in single and dKI mESCs (FXD repeats, *p* = 0.51; HD repeats, *p* = 0.60). Repeat PCR profiles of FXD (B) and HD (E) repeats in *Pms2*^+/+^ and *Pms2*^-/-^ dKI-mESCs carrying both FXD and HD repeats. The numbers in the day 42 profiles indicate the change in repeat number. The red dotted line indicates the starting allele. Changes in FXD (C) and HD (F) repeats number after 42 days in culture. The data represents the average of 3 technical replicates. The error bars indicate the standard deviations of the mean. Each dot represents one replicate.

We then integrated a doxycycline (DOX)-inducible construct expressing WT PMS2 into a *Pms2*^-*/*-^ dKI-mESC line and monitored the stability of both the FXD and the HD repeats over time in different concentrations of DOX. Without DOX treatment, small contractions like those seen in *Pms2*^-*/*-^ lines were also seen (Fig. 4). As the concentration of DOX was increased, so a progressive increase in repeat expansions was seen for both repeats with the highest level of expansion being seen at 30 ng/mL. The extent of expansions at this DOX concentration was comparable to the extent of expansions seen in *Pms2*^+1+^ cells. Higher concentrations of DOX resulted in a progressive decrease in expansions of both repeats. We repeated this experiment with a construct expressing PMS2 containing a D696N mutation in the nuclease domain (Fig. 5). In these lines, only small contractions, like those seen in the *Pms2*^-*/*-^ line, were seen for both repeats at all DOX concentrations tested (Fig. 5), including the DOX concentration that produced levels of PMS2 comparable to the levels seen in the WT parental line (Fig. 5D and Fig. S4). This suggests that an intact nuclease domain is required for PMS2’s promotion of expansion.

**Figure 4.**
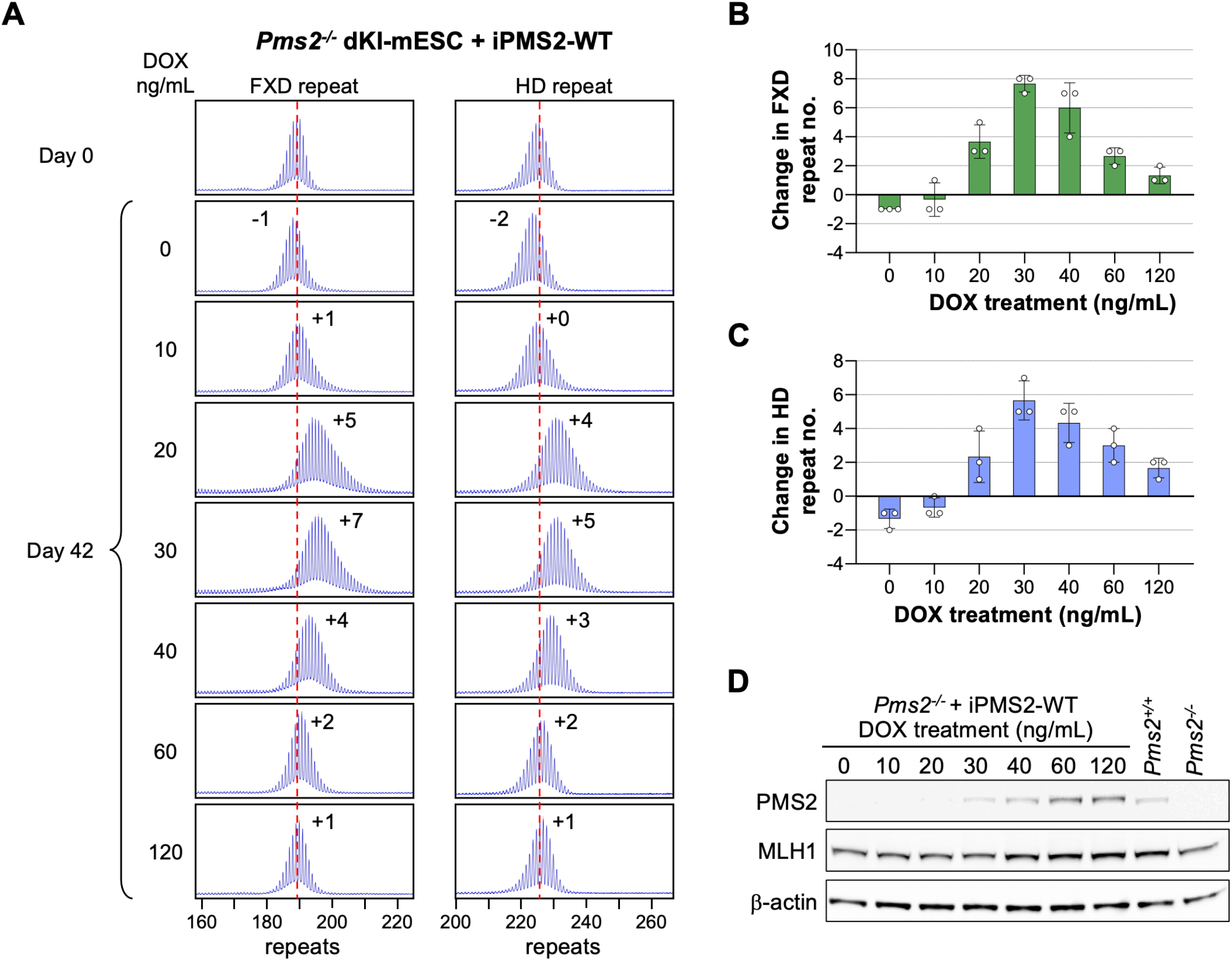
Effect of doxycycline-induced WT PMS2 expression on expansion of FXD and HD repeats in a double knock-in mESC model. (A) Repeat PCR profiles of FXD (left) and HD (right) repeats in *Pms2*^-/-^ dKI-mESCs expressing DOX-induced WT PMS2 (iPMS2-WT) at different concentrations of DOX after 42 days in culture. The number associated with each profile indicates the change in repeat number. The red dotted line indicates the starting allele. DOX concentrations producing similar levels of both DOX-induced WT and D696N versions of the DOX-induced PMS2 protein were used. Changes in FXD (B) and HD (C) repeat number at 42 days in *Pms2*^-/-^ dKI-mESCs expressing DOX-induced WT PMS2 at different concentrations of DOX. The data represents the average of 3 technical replicates. The error bars indicate the standard deviations of the mean. Each dot represents one replicate. (D) Western blots of whole-cell lysates from mESCs treated with the indicated concentrations of doxycycline and from *Pms2*^+/+^ and *Pms2*^-/-^ control mESCs. Blots were probed with antibodies indicated at left. The blots represent one of three technical replicates. Quantitative analysis for PMS2 and MLH1 are shown in Figure S4C. Representative examples of the full blots for binding of different antibodies are shown in Figure S7.

**Figure 5.**
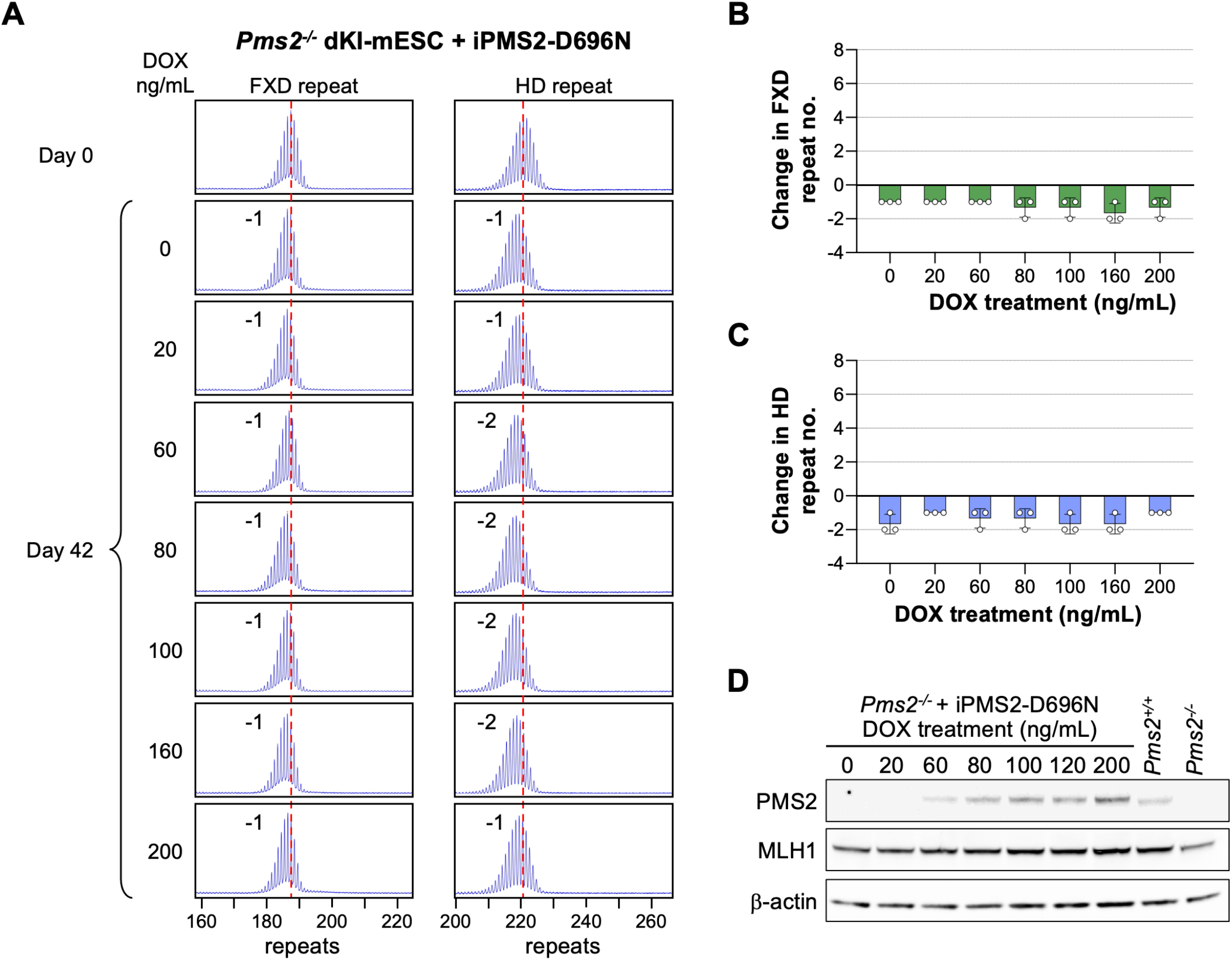
Effect of doxycycline-induced PMS2-D696N expression on expansion of FXD and HD repeats in a double knock-in mESC. (A) Repeat PCR profiles of FXD (left) and HD (right) repeats in *Pms2*^-/-^ dKI-mESCs expressing DOX-induced PMS2 D696N (iPMS2-D696N) at different concentrations of DOX after 42 days in culture. The number associated with each profile indicates the change in repeat number. The red dotted line indicates the starting allele. DOX concentrations producing similar levels of both DOX-induced WT and D696N versions of the DOX-induced PMS2 protein were used. Changes in FXD (B) and HD (C) repeat number at 42 days in *Pms2*^-/-^ dKI-mESCs expressing DOX-induced PMS2 D696N at different concentrations of DOX. The data represents the average of 3 technical replicates. The error bars indicate the standard deviations of the mean. Each dot represents one replicate. (D) Western blots of whole-cell lysates from mESCs treated with the indicated concentrations of doxycycline and from *Pms2*^+/+^ and *Pms2*^-/-^ control mESCs. Blots were probed with antibodies indicated at left. The blots represent one of three technical replicates. Quantitative analysis for PMS2 and MLH1 are shown in Figure S4D. Representative examples of the full blots for binding of different antibodies are shown in Figure S7.

## Discussion

We show here that in the same subset of tissues, including striatum and cortex, heterozygosity for *Pms2* resulted in a small, though not always significant, increase in expansion of both the FXD and HD repeats (Fig. 1 and Fig. S2 and Fig, S3). In nullizygous animals, these organs showed a further significant increase in expansion of both repeats. This would be consistent with PMS2 protecting against expansion. However, in other organs such as colon and testes, nullizygosity for *Pms2* resulted in the loss of most, if not all, expansions in both sets of mice, consistent with the requirement of PMS2 for expansion. This is very different from what is seen in *Mlh3* null animals where no expansions of either the HD or FXD repeat was seen in any tissue (16, 17). Counterintuitively, some of the organs in which PMS2 is required for expansion showed more expansion in heterozygous animals than in WT or null animals. This may be related to the fact that while heterozygosity for *Pms2* was associated with a ∼50% reduction in PMS2 protein in organs like testes, a decline in MLH1 levels was only seen in *Pms2* null animals (Fig. S1). Since PMS1 and PMS2 compete for binding to MLH1 with binding being important for their stability (34), the increase in expansions in some organs of heterozygous animals may reflect an increase in the amount of PMS1 able to bind MLH1. A role for PMS2 in driving expansions is supported by our demonstration that in *Pms2*^-*/*-^ mESCs no expansions of either the FXD or the HD repeats were seen (Fig. 3); but that when levels of PMS2 are systematically increased, expansion rates rise, peaking at rates similar to those seen in WT cells for both repeats (Fig. 4). The subsequent decrease in expansions that were seen with higher levels of DOX may reflect competition between PMS2 and PMS1 for the available MLH1.

Therefore, the different effects of PMS2 previously reported in different model systems of different REDs likely do not reflect fundamentally different mechanisms of instability in these diseases, but rather the ability of PMS2 to promote or repress expansion in different tissues or cell types. The fact that a PMS2 expression construct with a point mutation in the nuclease domain is unable to restore expansions (Fig. 5) suggests that PMS2’s nuclease activity is required for its role in promoting expansions as we had previously shown for MLH3 (20).

How PMS2 can both promote and protect against expansions is an open question. Our demonstration of the requirement for both the MLH3 and the PMS2 nuclease domains for expansion in mESCs suggests that both proteins can contribute to the generation of expansions and do so via their ability to nick the expansion substrate. This is consistent with current models for expansion in which the expansion substrate has two loop-outs (31). Since MLH3 is required for all expansions while PMS2 is not, one explanation may be that PMS2 is able to complement MLH3’s processing of the expansion substrate to generate expansions, but is unable to act alone to do so. This may be related to previous work showing that *in vitro* MLH3 cleaves the DNA strand opposite any loop-out to which it binds (35), while on nicked substrates PMS2 cleaves the nicked strand and in the absence of a nick, has an equal probability of cutting either strand (22, 36, 37). We hypothesize that expansions result when both loop-outs are processed via cleavage that occurs on the strand opposite each loop-out. Such an intermediate would then be processed via gap repair such that both sets of the loop-out bases would be incorporated into the “repaired” strands as illustrated in Fig. S5 (i). If MutLγ cleavage *in vivo* occurs as it does *in vitro*, then MutLγ processing of both loop-outs would generate such an intermediate. MutLα cleavage on the other hand would generate a wider range of intermediates, most of which would not result in the incorporation of both loop-outs. While at some frequency a combination of both MutLγ and MutLα cleavage would be predicted to generate an intermediate that could result in expansion, MutLα processing of both loop-outs would not do so since a nick at one loop-out would favor a nick on the same strand at the other. In this scenario, PMS2 protects against expansion by favoring the error-free repair of the expansion substrate. However, in some situations it acts together with MLH3 to allow misincorporation of the loop-outs, thereby generating expansions. Why some cell populations require MutLα for expansions while others do not remains an open question. It may be related to the relative abundance of the other factors that contribute to the expansion process. However, since expansion only occurs in a subset of cells in many tissues *e*.*g*., the stem cells of the testes or small intestine in FXD mice (38) and the medium spiny neurons and CHAT+ interneurons in the striatum of HD patients (39, 40), properly addressing this question would require proteomic analysis of low abundance proteins in small cell populations.

Nevertheless, our data demonstrate that PMS2 acts on expansions of both the HD and FXD repeats in a similar way, both contributing to their production in some cases and protecting against them in others. This finding lends support to the idea that different REDs share a similar or common expansion mechanism. This has implications for our understanding of the mechanisms controlling repeat instability in this group of disorders. It also increases the confidence that a successful approach for reducing somatic expansions in one of these diseases might be useful to the other diseases in this group as well.

## Materials and Methods

### Reagents and services

Reagents were from Sigma-Aldrich (St Louis, MO) unless otherwise stated. Primers were from Life Technologies (Grand Island, NY). Capillary electrophoresis of fluorescently labeled PCR products was carried out by the Roy J Carver Biotechnology Center, University of Illinois (Urbana, IL) and Psomagen (Rockville, MD).

### Mouse generation, breeding, and maintenance

Embryos of *Pms2* mutant mice (41) were obtained from The Jackson Laboratory (Bar Harbor, ME; JAX stock #010945) and recovered by NIDDK Laboratory Animal Sciences section (LASS) using standard procedures. The HD mice (zQ175: B6J.129S1-*Htt*^*tm1Mfc*^190ChdiJ) (42, 43) were acquired from The Jackson Laboratory (Bar Harbor, ME; JAX stock #027410). The FXD mice containing expanded CGG-repeats inserted into the endogenouse mouse *Fmr1* locus (Fmr1^tm1Usdn^; MGI:3711215) (44) have been previously described. *Pms2* mutant mice were crossed to FXD and HD mice to generate animals that were heterozygous for *Pms2*. These mice were then crossed again with FXD or HD mice to generate mice homozygous for the *Pms2* mutation. All mice were on a C57BL/6J background. Mice were maintained in a manner consistent with the Guide for the Care and Use of Laboratory Animals (NIH publications no. 85-23, revised 1996) and in accordance with the guidelines of the NIDDK Animal Care and Use Committee, who approved this research (ASP-K021-LMCB-21).

### Generation of doxycycline-inducible Pms2 constructs

Two plasmids, iPMS2-WT and iPMS2-D696N, were generated to express either WT PMS2 or a nuclease-dead version of PMS2 (D696N) (22) under the control of a doxycycline-inducible promoter. In both of these constructs, an hPGK and an mPGK promoter drive constitutive expression of the doxycycline-responsive TetOn-3G gene and the mClover3 green fluorescent reporter gene (from pKK-TEV-mClover3, Addgene #105795), respectively, as shown in Fig. S5A. The *Pms2* coding sequence was placed downstream of the doxycycline-inducible promoter. The sequence encoding PMS2 corresponds to NCBI Reference Sequence NP_032912.2, with a 1x FLAG epitope sequence inserted immediately after the first codon, and the final codon replaced with an alternate stop codon. In the PMS2-D696N version of the construct, an AAC codon (asparagine) replaces the GAC codon (aspartic acid) at the position corresponding to amino acid 696 of the WT PMS2. These elements are flanked by left and right ROSA homology arms from pROSA26-1 (Addgene #21714) for targeting the construct to endogenous ROSA26 locus. Fragments were combined using standard techniques including Gibson Assembly and NEBuilder HiFi reagents (New England Biolabs, Ipswich, MA). Final construct sequences were confirmed by Sanger sequencing (Psomagen, Inc., Rockville, MD) and whole-plasmid sequencing (Plasmidsaurus, Inc., Louisville, KY).

### Generation and culture of mESCs

The double knock-in mouse ESC (dKI-mESC) carrying both FXD and HD knock-in alleles were derived from embryos obtained by crossing FXD and HD mice using standard procedures and routinely cultured as previously described (45). *Pms2* null alleles were generated in an dKI-mESC line with ∼190 FXD repeats and ∼228 HD repeats using a CRISPR-Cas9 approach as described previously (23). A *Pms2*^-/-^ dKI-mESC line was transfected with constructs that express either WT PMS2 (iPMS2-WT) or a nuclease-dead version of PMS2 (iPMS2-D696N) under the control of a doxycycline-inducible promoter, described above. These constructs were targeted to the ROSA26 locus of the *Pms2*^-*/*-^ dKI-mESC lines by co-transfection with a Cas9-expressing plasmid (14), that had been modified to contain gRNAs for the ROSA26 locus. Single-cell-derived lines with stable integration of the transfected construct were identified by expression of a constitutively expressed mClover3 fluorescent reporter protein. Culture media for mESCs was supplemented with DOX at concentrations indicated for various experiments. DOX-induction of the WT and D696N PMS2 was verified both by RT-qPCR and western blotting using standard procedures. For a given DOX concentration, the amount of DOX-induced WT PMS2 protein produced was ∼2-fold higher than the D696N protein (Fig. S4C and D). However, this does not reflect differences in the protein stability since the amount of PMS2 mRNA produced showed a similar difference (Fig. S4B). The reason for this difference is unclear but is frequently seen with this integration strategy and may reflect differences in the number of copies of the expression construct that were integrated.

### Western bloting of mESC

Cells for western blotting were cultured in 6-well plates for 2 days with daily medium changes. Cells were rinsed with DPBS (Dulbecco’s phosphate buffered saline) and treated with 1x TrypLE Select (ThermoFisher Scientific, Waltham, MA) in DPBS. Once cells detached from the plate, 15% tetracyline-free serum in cell culture medium was added and cells were dissociated by pipetting. Dissociated cells were pelleted, washed in cold DPBS with protease inhibitors, re pelleted, and frozen. Cell pellets were lysed in T-PER Tissue Protein Extraction Reagent (ThermoFisher Scientific). The protein concentrations were determined using a Bio-Rad protein assay kit (Bio-Rad, Hercules, CA). For each sample, 12 µg protein per lane was analyzed on 4-12% bis-tris gels in MOPS running buffer (ThermoFisher Scientific). Proteins were transferred to nitrocellulose membranes using Trans-Blot Turbo transfer buffer (Bio-Rad Laboratories, Hercules, CA). Antibodies were diluted in 5% ECL Prime Blocking Agent (Cytiva, Wilmington, DE) in tris-buffered saline: anti-PMS2 (1:1000; clone A16-4, 556415, BD Pharmingen), anti-MLH1 (1:10000, ab92312, Abcam), anti-β-actin (1:15000, 15G5A11/E2, MA1-140, ThermoFisher Scientific), goat anti-mouse lgG:Dylight 800 (1:2500, Bio-Rad Laboratories), goat anti-rabbit StarBright Blue 520 (1:2500, Bio-Rad Laboratories). The band intensity was determined using Bio-Rad Image Lab v 6.1.0 build 7 software. The protein levels were normalized to β-actin and quantitative analysis results are shown as the fold change relative to the level in *Pms2*^+*/*+^cells.

### DNA isolation

DNA for genotyping was extracted from mouse tails collected at 3-weeks-old, or weaning, using KAPA Mouse Genotyping Kit (KAPA Biosystems, Wilmington, MA). DNA was isolated from a variety of tissues that were collected from 4- and 8-month-old male mice using a Maxwell®16 Mouse Tail DNA Purification kit (Promega, Madison, WI) according to the manufacturer’s instructions. A 5 cm section of the jejunum was collected as the small intestine sample and a 5 cm distal colon sample was collected upstream of the anus as previously described (46). Sperm collection and DNA preparation were as previously described (47). DNA was purified from mESCs as described previously (23).

### Genotyping and analysis of repeat number

Genotyping of *Pms2* was carried out using the KAPA mouse genotyping kit (KAPA Biosystems) according to manufacturer’s instructions with primers JAX-9366 (5’-TTCGGTGACAGATTTGTAAATG-3’) and JAX-9367 (5’-TCACCATAAAAATAGTTTCCCG-3’) used to detect the WT *Pms2* allele and JAX-9366 and JAX-9368 (5’-TTTACGGAGCCCTGGC-3’) to detect the mutant *Pms2* allele. The PCR mix for the *Pms2* allele contained 2 µL template DNA, 1X KAPA2G Fast HotStart Genotyping Mix (KAPA Biosystems, Wilmington, MA), and 0.5 µM each of the primers. The *Pms2* allele PCR conditions were 95°C for 3 minutes; 35 cycles of 95°C for 15 seconds, 60°C for 15 seconds and 72°C for 15 seconds; followed by 72°C for 3 minutes. Genotyping and repeat size analysis of the *Fmr1* and *Htt* alleles was performed using a fluorescent PCR assay with fluorescein amidite (FAM)-labeled primer pairs. The primers FAM labeled FraxM4 (FAM-5’-CTTGAGGCCCAGCCGCCGTCGGCC-3’) and FraxM5 (5’ CGGGGGGCGTGCGGTAACGGCCCAA-3’) were used for the *Fmr1* allele (44). The PCR mix for *Fmr1* allele contained 3 µL (150 ng) template DNA, 1X KAPA2G Fast HotStart Genotyping Mix, 2.4 M betaine, 2% DMSO, 0.5 µM each of the primers and additional of 125 µM each of dCTP and dGTP. The PCR cycling parameters for the *Fmr1* allele were 95°C for 10 minutes; 35 cycles of 95°C for 30 seconds, 65°C for 30 seconds and 72°C for 90 seconds; followed by 72°C for 10 minutes. The primers FAM-labeled HU3 (FAM-5’-GGCGGCTGAGGAAGCTGAGGA-3’) and Htt EX1-F1 (5’-GCAACCCTGGAAAAGCTGATGAAGGC-3’) were used for the *Htt* allele. The PCR mix for the *Htt* allele contained 2 µL (100 ng) DNA template, 1x KAPA2G Fast HotStart Genotyping Mix, 1.2 M betaine, 1% DMSO, and 0.5 µM each of the primers. The *Htt* allele was amplified by touchdown PCR using the following parameters: 95°C for 10 minutes; 10 cycles of 95°C for 30 seconds, 72°C with -1°C/cycle for 30 seconds and 72°C for 90 seconds; 28 cycles of 95°C for 30 seconds, 63°C for 30 seconds and 72°C for 90 seconds; followed by 72°C for 10 minutes. The *Fmr1* and *Htt* PCR products were resolved by capillary electrophoresis on an ABI Genetic Analyzer and the resultant fsa files were displayed using a previously described custom R script (48) that is available upon request. The tail sample that was taken at 3-weeks or weaning was used as a proxy-indicator of the original inherited allele size. The expansion index (EI) was calculated in the same way as the somatic instability index (49), but only peaks larger than the original inherited allele were considered, with a cutoff of 10% relative peak height threshold. The repeat number changes were determined by subtracting the number of repeats in the modal allele from the number of repeats in the original inherited allele.

### Statistical analyses

Statistical analyses were performed using GraphPad Prism 10.2. For comparisons of EI or repeat number changes in samples with different genotypes or ages, statistical significance was assessed using the two-way ANOVA with Tukey’s multiple comparisons correction. Linear regression was performed to model the relationship between the repeat number changes and the days in culture, with the slope representing the rate of change of the repeats with respect to the day. The expansion rate per week was calculated by multiplying the expansion rate per day by 7. The repeat rate in single and double knock-in mESC were compared using a two-tailed unpaired t-tests.

## Acknowledgements

This research was supported by the Intramural Research Program of the NIH, The National Institute of Diabetes and Digestive and Kidney Diseases (NIDDK) and CHDI foundation. The authors want to thank Dr. Farid Kadyrov, Dr. Darren Monckton, Dr. Jasmine Donaldson, Dr. Gabriel Balmus for helpful discussions, and the members of the Usdin laboratory for the helpful comments and suggestions. We would like to acknowledge all the hard work done by the staff who take care of our mice and without whom this work would not have been possible.

## Author contributions

Conceptualization: C.J.M., K.U., X.Z.; Methodology: C.J.M., B.E.H., H.L., X.Z.; Validation: C.J.M., X.Z.; Formal analysis: D.A.J., C.J.M., A.W., B.E.H., H.L., X.Z.; Investigation: D.A.J., C.J.M., A.W., K.A., X.Z.; Resources: K.U.; Data curation: D.A.J., C.J.M., A.W., X.Z.; Writing - original draft: K.U., X.Z.; Writing - review & editing: D.A.J., C.J.M., A.W., K.U., X.Z.; Visualization: C.J.M., B.E.H., H.L., X.Z.; Supervision: K.U., X.Z.; Project administration: X.Z.; Funding acquisition: K.U.

## Data availability

All data generated or analyzed during this study are included in this published article and its Supplementary Information files.

## References

1. C. Depienne, J. L. Mandel, 30 years of repeat expansion disorders: What have we learned and what are the remaining challenges? Am J Hum Genet 108, 764–785 (2021).

2. K. Ibanez et al., Increased frequency of repeat expansion mutations across different populations. Nat Med 30, 3357–3368 (2024).

3. Genetic Modifiers of Huntington’s Disease (GeM_HD) Consortium, Identification of Genetic Factors that Modify Clinical Onset of Huntington’s Disease. Cell 162, 516–526 (2015).

4. C. Bettencourt et al., DNA repair pathways underlie a common genetic mechanism modulating onset in polyglutamine diseases. Ann Neurol 79, 983–990 (2016).

5. M. Flower et al., MSH3 modifies somatic instability and disease severity in Huntington’s and myotonic dystrophy type 1. Brain 142, 1876–1886 (2019).

6. Genetic Modifiers of Huntington’s Disease (GeM_HD) Consortium, CAG Repeat Not Polyglutamine Length Determines Timing of Huntington’s Disease Onset. Cell 178, 887–900 e814 (2019).

7. Y. H. Hwang et al., Both cis and trans-acting genetic factors drive somatic instability in female carriers of the FMR1 premutation. Sci Rep 12, 10419 (2022).

8. R. Wiggins, A. Feigin, Emerging therapeutics in Huntington’s disease. Expert Opin Emerg Drugs 26, 295–302 (2021).

9. W. J. van den Broek et al., Somatic expansion behaviour of the (CTG)n repeat in myotonic dystrophy knock-in mice is differentially affected by Msh3 and Msh6 mismatch-repair proteins. Hum Mol Genet 11, 191–198 (2002).

10. L. Foiry et al., Msh3 is a limiting factor in the formation of intergenerational CTG expansions in DM1 transgenic mice. Hum Genet 119, 520–526 (2006).

11. S. Tome et al., MSH3 polymorphisms and protein levels affect CAG repeat instability in Huntington’s disease mice. PLoS Genet 9, e1003280 (2013).

12. X. N. Zhao et al., Mutsbeta generates both expansions and contractions in a mouse model of the Fragile X-associated disorders. Hum Mol Genet 24, 7087–7096 (2015).

13. J. C. L. Roy et al., Somatic CAG expansion in Huntington’s disease is dependent on the MLH3 endonuclease domain, which can be excluded via splice redirection. Nucleic Acids Res 49, 3907–3918 (2021).

14. B. Hayward et al., All three MutL complexes are required for repeat expansion in a human stem cell model of CAG-repeat expansion mediated glutaminase deficiency. Sci Rep 14, 13772 (2024).

15. R. Ferguson, R. Goold, L. Coupland, M. Flower, S. J. Tabrizi, Therapeutic validation of MMR-associated genetic modifiers in a human ex vivo model of Huntington disease. Am J Hum Genet 111, 1165–1183 (2024).

16. R. M. Pinto et al., Mismatch repair genes Mlh1 and Mlh3 modify CAG instability in Huntington’s disease mice: genome-wide and candidate approaches. PLoS Genet 9, e1003930 (2013).

17. X. Zhao, Y. Zhang, K. Wilkins, W. Edelmann, K. Usdin, MutLgamma promotes repeat expansion in a Fragile X mouse model while EXO1 is protective. PLoS Genet 14, e1007719 (2018).

18. A. Halabi, K. T. B. Fuselier, E. Grabczyk, GAA*TTC repeat expansion in human cells is mediated by mismatch repair complex MutLgamma and depends upon the endonuclease domain in MLH3 isoform one. Nucleic Acids Res 46, 4022–4032 (2018).

19. E. Cannavo et al., Expression of the MutL homologue hMLH3 in human cells and its role in DNA mismatch repair. Cancer Res 65, 10759–10766 (2005).

20. B. E. Hayward, P. J. Steinbach, K. Usdin, A point mutation in the nuclease domain of MLH3 eliminates repeat expansions in a mouse stem cell model of the Fragile X-related disorders. Nucleic Acids Res 48, 7856–7863 (2020).

21. F. Palombo et al., hMutSbeta, a heterodimer of hMSH2 and hMSH3, binds to insertion/deletion loops in DNA. Curr Biol 6, 1181–1184 (1996).

22. F. A. Kadyrov, L. Dzantiev, N. Constantin, P. Modrich, Endonucleolytic function of MutLαlpha in human mismatch repair. Cell 126, 297–308 (2006).

23. C. J. Miller, G. Y. Kim, X. Zhao, K. Usdin, All three mammalian MutL complexes are required for repeat expansion in a mouse cell model of the Fragile X-related disorders. PLoS Genet 16, e1008902 (2020).

24. R. Mouro Pinto et al., In vivo CRISPR-Cas9 genome editing in mice identifies genetic modifiers of somatic CAG repeat instability in Huntington’s disease. Nat Genet 57, 314– 322 (2025).

25. N. Wang et al., Distinct mismatch-repair complex genes set neuronal CAG-repeat expansion rate to drive selective pathogenesis in HD mice. Cell 188, 1524–1544 e1522 (2025).

26. L. Y. Kadyrova, F. F. Kadyrov, B. Hayward, K. Usdin, F. A. Kadyrov, Mechanism of MutLβ-dependent DNA expansions. Proc Natl Acad Sci U S A In Press (2026).

27. R. Mouro Pinto et al., Identification of genetic modifiers of Huntington’s disease somatic CAG repeat instability by in vivo CRISPR-Cas9 genome editing. bioRxiv 10.1101/2024.06.08.597823 (2024).

28. R. L. Bourn et al., Pms2 suppresses large expansions of the (GAA.TTC)n sequence in neuronal tissues. PLoS One 7, e47085 (2012).

29. V. Campuzano et al., Friedreich’s ataxia: autosomal recessive disease caused by an intronic GAA triplet repeat expansion. Science 271, 1423–1427 (1996).

30. H. G. Harley et al., Unstable DNA sequence in myotonic dystrophy. Lancet 339, 1125– 1128 (1992).

31. M. Gomes-Pereira, M. T. Fortune, L. Ingram, J. P. McAbney, D. G. Monckton, Pms2 is a genetic enhancer of trinucleotide CAG.CTG repeat somatic mosaicism: implications for the mechanism of triplet repeat expansion. Hum Mol Genet 13, 1815–1825 (2004).

32. R. Lozano, C. A. Rosero, R. J. Hagerman, Fragile X spectrum disorders. Intractable Rare Dis Res 3, 134–146 (2014).

33. A. B. P. van Kuilenburg et al., Glutaminase Deficiency Caused by Short Tandem Repeat Expansion in GLS. N Engl J Med 380, 1433–1441 (2019).

34. M. Raschle, G. Marra, M. Nystrom-Lahti, P. Schar, J. Jiricny, Identification of hMutLbeta, a heterodimer of hMLH1 and hPMS1. J Biol Chem 274, 32368–32375 (1999).

35. L. Y. Kadyrova, V. Gujar, V. Burdett, P. L. Modrich, F. A. Kadyrov, Human MutLgamma, the MLH1-MLH3 heterodimer, is an endonuclease that promotes DNA expansion. Proc Natl Acad Sci U S A 117, 3535–3542 (2020).

36. A. Pluciennik et al., PCNA function in the activation and strand direction of MutLαlpha endonuclease in mismatch repair. Proc Natl Acad Sci U S A 107, 16066–16071 (2010).

37. A. Pluciennik et al., Extrahelical (CAG)/(CTG) triplet repeat elements support proliferating cell nuclear antigen loading and MutLαlpha endonuclease activation. Proc Natl Acad Sci U S A 110, 12277–12282 (2013).

38. X. N. Zhao, K. Usdin, Timing of Expansion of Fragile X Premutation Alleles During Intergenerational Transmission in a Mouse Model of the Fragile X-Related Disorders. Front Genet 9, 314 (2018).

39. K. Matlik et al., Cell-type-specific CAG repeat expansions and toxicity of mutant Huntingtin in human striatum and cerebellum. Nat Genet 56, 383–394 (2024).

40. R. E. Handsaker et al., Long somatic DNA-repeat expansion drives neurodegeneration in Huntington’s disease. Cell 188, 623–639 e619 (2025).

41. S. M. Baker et al., Male mice defective in the DNA mismatch repair gene PMS2 exhibit abnormal chromosome synapsis in meiosis. Cell 82, 309–319 (1995).

42. L. B. Menalled, J. D. Sison, I. Dragatsis, S. Zeitlin, M. F. Chesselet, Time course of early motor and neuropathological anomalies in a knock-in mouse model of Huntington’s disease with 140 CAG repeats. J Comp Neurol 465, 11–26 (2003).

43. J. C. Grima et al., Mutant Huntingtin Disrupts the Nuclear Pore Complex. Neuron 94, 93–107 e106 (2017).

44. A. Entezam et al., Regional FMRP deficits and large repeat expansions into the full mutation range in a new Fragile X premutation mouse model. Gene 395, 125–134 (2007).

45. I. Gazy, C. J. Miller, G. Y. Kim, K. Usdin, CGG Repeat Expansion, and Elevated Fmr1 Transcription and Mitochondrial Copy Number in a New Fragile X PM Mouse Embryonic Stem Cell Model. Front Cell Dev Biol 8, 482 (2020).

46. X. Zhao et al., Stool is a sensitive and noninvasive source of DNA for monitoring expansion in repeat expansion disease mouse models. Dis Model Mech 15 (2022).

47. X. Zhao, H. Lu, P. K. Dagur, K. Usdin, Isolation and Analysis of the CGG-Repeat Size in Male and Female Gametes from a Fragile X Mouse Model. Methods Mol Biol 2056, 173–186 (2020).

48. B. E. Hayward, Y. Zhou, D. Kumari, K. Usdin, A Set of Assays for the Comprehensive Analysis of FMR1 Alleles in the Fragile X-Related Disorders. J Mol Diagn 18, 762–774 (2016).

49. J. M. Lee et al., A novel approach to investigate tissue-specific trinucleotide repeat instability. BMC Syst Biol 4, 29 (2010).

