## Supplemental data for "Reconciling the effects of PMS2 in different repeat expansion disease models supports a common expansion mechanism"

### Supplementary Information

#### Supplemental Methods

*RT-qPCR*

Confluent mESCs were diluted 1:6, plated into 12-well tissue culture dishes and grown for 2 days with a medium change after 1 day. Total RNA was prepared using the RNeasy Mini Kit according to the manufacturer's directions (QIAGEN, Germantown, MD) and genomic DNA was removed by on-column digestion with DNAase I (New England Biolabs, Ipswich, MA). Reverse transcription was performed using SuperScript IV VILO Master Mix (Thermo Fisher Scientific, Waltham, MA) as directed by the manufacturer. Quantitative PCR was performed using PowerUp SYBR Green Master Mix (ThermoFisher Scientific) with KiCqStart SYBR Green Primers (Millipore-Sigma, Burlington, MA) targeting mouse *Pms2* (FM2_Pms2, 5'-GTTTCCGTTGACTCAGAATG-3' and RM2_Pms2, 5'-GCAGTATGCAGCTTTACTAAG-3') and targeting *Actb* as a reference gene (FM1_Actb, 5'-GATGTATGAAGGCTTTGGTC-3' and RM1_Actb, 5'-TGTGCACTTTTATTGGTCTC-3'). PCR conditions: 50°C for 2 minutes; 95°C for 2 minutes; 40 cycles of 95°C for 2 seconds, 57°C for 15 seconds, 72°C for 30 seconds; and followed by melt-curve analysis to assess the fidelity of the reactions.

*Western blotting of mouse tissue*

The flash frozen testes and striatum samples were collected from 4-month-old mice. Total proteins were extracted using the T-PER protein extraction reagent (Pierce Biotechnology, Rockford, IL) supplemented with complete, Mini, EDTA-free protease inhibitor cocktail (Roche

Applied Science, Indianapolis, IN) according to the manufacturer's instructions. Briefly, tissues were weight and homogenized using a tissue homogenizer (Precellys 24, Bertin Technologies, Berlin, Germany) with T-PER reagent. Tissue debris were discarded after centrifuge at 10,000 g for 5 minutes. Supernatant were collected and the protein concentrations were determined using a Bio-Rad protein assay kit (Bio-Rad, Hercules, CA). After adding the LDS-Sample Buffer (Thermo Fisher Scientific) and Sample Reducing agent (Thermo Fisher Scientific), proteins were heated for 10 minutes at 70°C. A total of 50 ug lysate was resolved by electrophoresis on 4–12% NuPAGE Novex Tris-Bis gels (Thermo Fisher Scientific) in 1x MOPS SDS Running buffer (Thermo Fisher Scientific) and transferred to nitrocellulose membranes using the Trans-Blot Turbo RTA transfer kit (Bio-Rad) according to the manufacturer's instructions. Antibodies were diluted in 5% ECL Prime Blocking Agent (Cytiva, Wilmington, DE) in tris-buffered saline. The following primary antibodies were used for immunoblotting: anti-PMS2 (1:1000; clone A16-4, 556415, BD Pharmingen), anti-MLH1 (1:5000, ab92312, Abcam), anti-β-actin (1:10000, 15G5A11/E2, MA1-140, Thermo Fisher Scientific). The following secondary antibodies were used: anti-mouse IgG (1:5000, 12-349, Millipore), and anti-rabbit IgG (1:5000, GENA934, Millipore-Sigma). The band intensity was determined using ImageJ 2 software. The protein levels were normalized to β-actin and quantitative analysis results are shown as the fold change relative to the level in *Pms2*$^{+/+}$ mice.

### Supplemental figures and legends

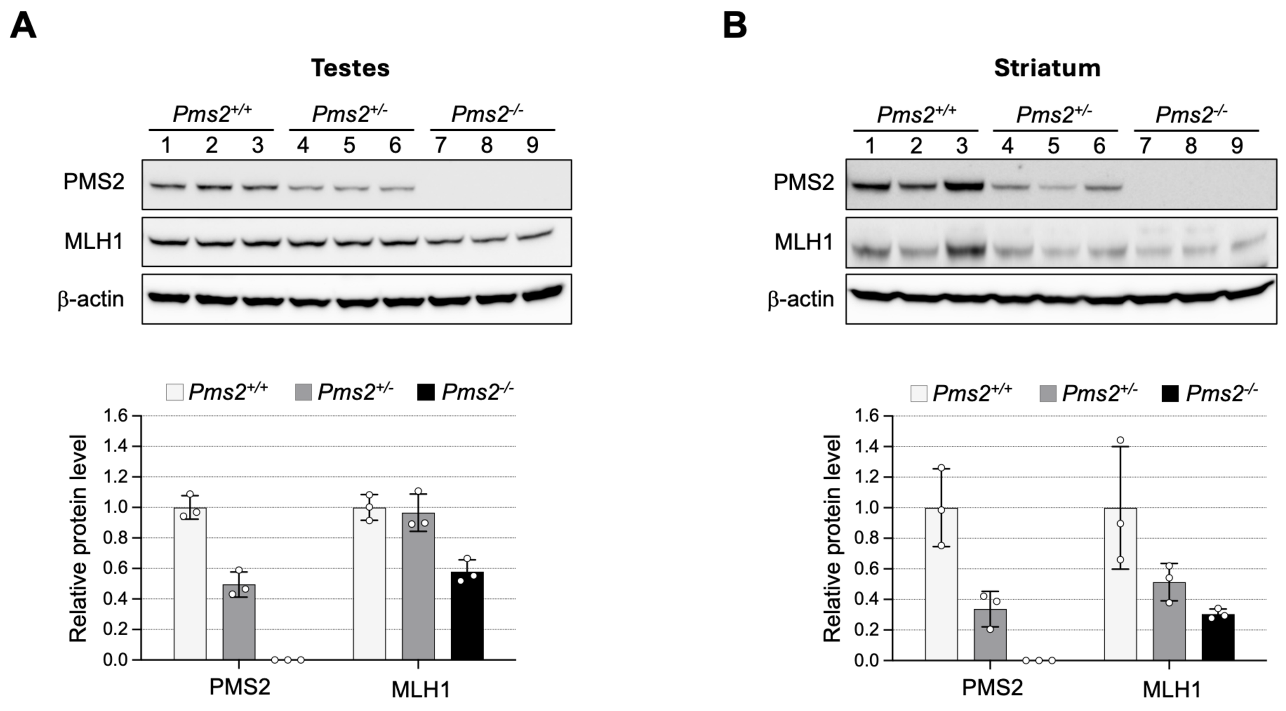


**Figure S1. The levels of PMS2 protein in mouse testes and striatum.** Western blot of testes (A) and striatum (B) samples from three 4-month-old *PMS2*$^{+/+}$, *PMS2*$^{+/-}$, and *PMS2*$^{-/-}$ animals. Blots were probed with the antibodies indicated. The uncropped blots showing the binding of antibodies to PMS2, MLH1, and b-actin are shown in Fig. S7. The band intensity was determined using ImageJ 2 software as described in the Materials and Methods. The PMS2 and MLH1 protein levels were normalized to β-actin. The bar graph at the bottom of the figure shows the fold change relative to the level in *Pms2*$^{+/+}$ mice. The data represents the average of 3 biological replicates. The error bars indicate the standard deviations of the mean. Each dot represents one replicate.

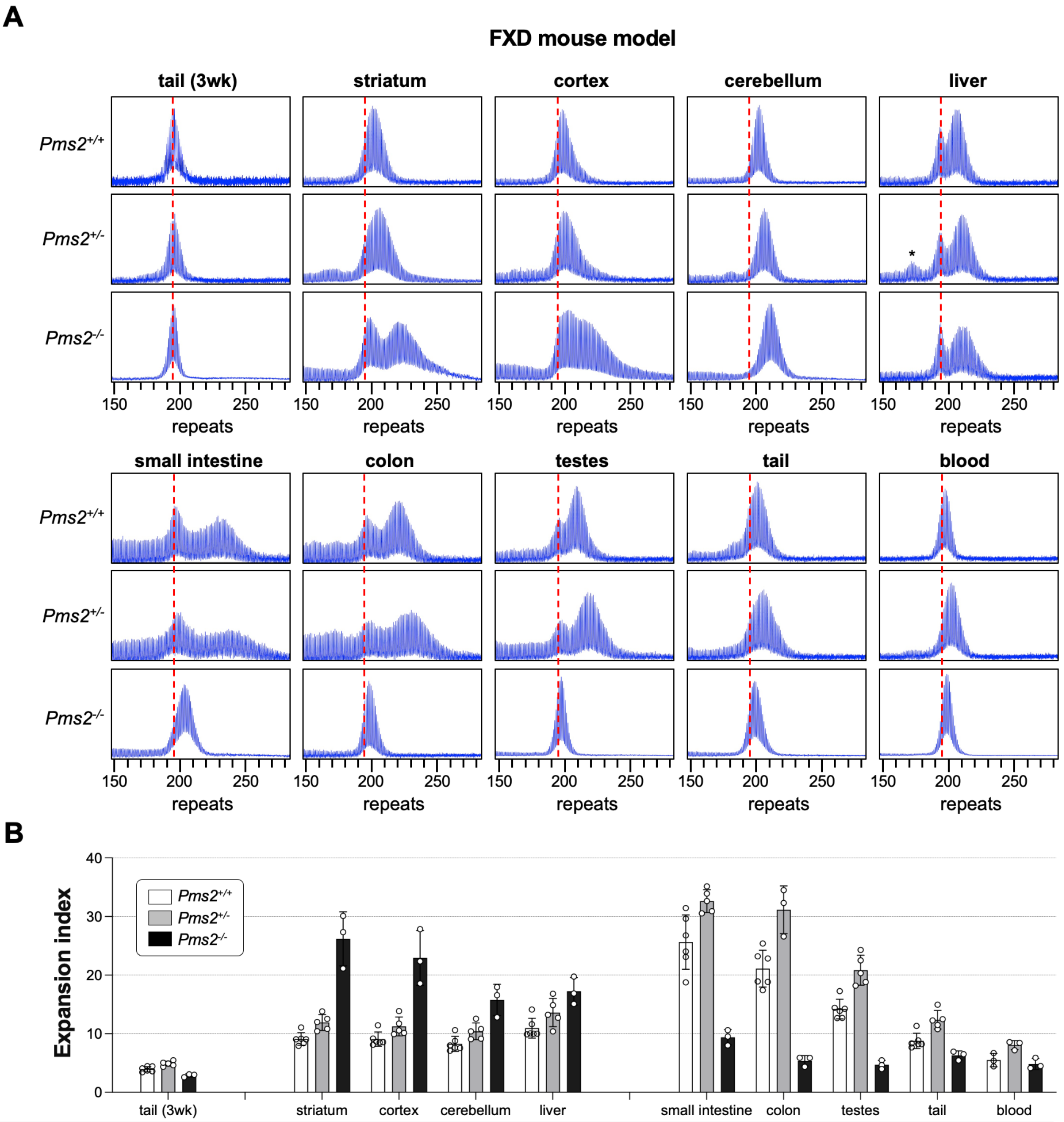


| | tail (3wk) | | striatum | cortex | cerebellum | liver | | small intestine | colon | testes | tail | blood |
|---|---|---|---|---|---|---|---|---|---|---|---|---|
| ***Pms2*^+/+^ vs *Pms2*^+/-^** | 0.7541 | | 0.0815 | 0.2224 | 0.2349 | 0.1067 | | **<0.0001** | **<0.0001** | **<0.0001** | **0.0178** | 0.3145 |
| ***Pms2*^+/+^ vs *Pms2*^-/-^** | 0.7618 | | **<0.0001** | **<0.0001** | **<0.0001** | **0.0002** | | **<0.0001** | **<0.0001** | **<0.0001** | 0.2371 | 0.9252 |
| ***Pms2*^+/-^ vs *Pms2*^-/-^** | 0.412 | | **<0.0001** | **<0.0001** | **0.0025** | 0.0578 | | **<0.0001** | **<0.0001** | **<0.0001** | **0.0005** | 0.1632 |

**Figure S2. The effect of *Pms2* deficiency on repeat expansion in different tissues of an FXD mouse model.** (A) Representative repeat PCR profiles from tail DNA taken at 3 weeks (3 wk) and different organs of 4-month-old *Pms2*$^{+/+}$, *Pms2*$^{+/-}$ and *Pms2*$^{-/-}$ FXD male mice with 196 repeats. The dashed lines represent the sizes of the original inherited alleles as ascertained from the tail DNA taken at 3 weeks. (B) Comparison of the expansion index (EI) in the indicated organs of 4-month-old *Pms2*$^{+/+}$, *Pms2*$^{+/-}$ and *Pms2*$^{-/-}$ FXD mice with an average of 194 repeats in the original allele. The colon data represent the average of 6 *Pms2*$^{+/+}$, 3 *Pms2*$^{+/-}$ and 3 *Pms2*$^{-/-}$ mice with 185-210 repeats. The blood data represent the average of 3 *Pms2*$^{+/+}$, 3 *Pms2*$^{+/-}$ and 3 *Pms2*$^{-/-}$ mice in the same repeat range. The data from other organs represents the average of 6 *Pms2*$^{+/+}$, 5 *Pms2*$^{+/-}$ and 3 *Pms2*$^{-/-}$ mice in the same repeat range. The error bars indicate the standard deviations of the mean. Each dot represents one animal. In each organ, the EIs for different genotypes were compared using a two-way ANOVA with correction for multiple testing as described in the Materials and Methods. The adjusted *p*-values are listed in the table below. The asterisks in the *Pms2*$^{+/-}$ liver sample indicates a contracted allele that is also present in other organs and not a specific contraction caused by PMS2 deficiency.

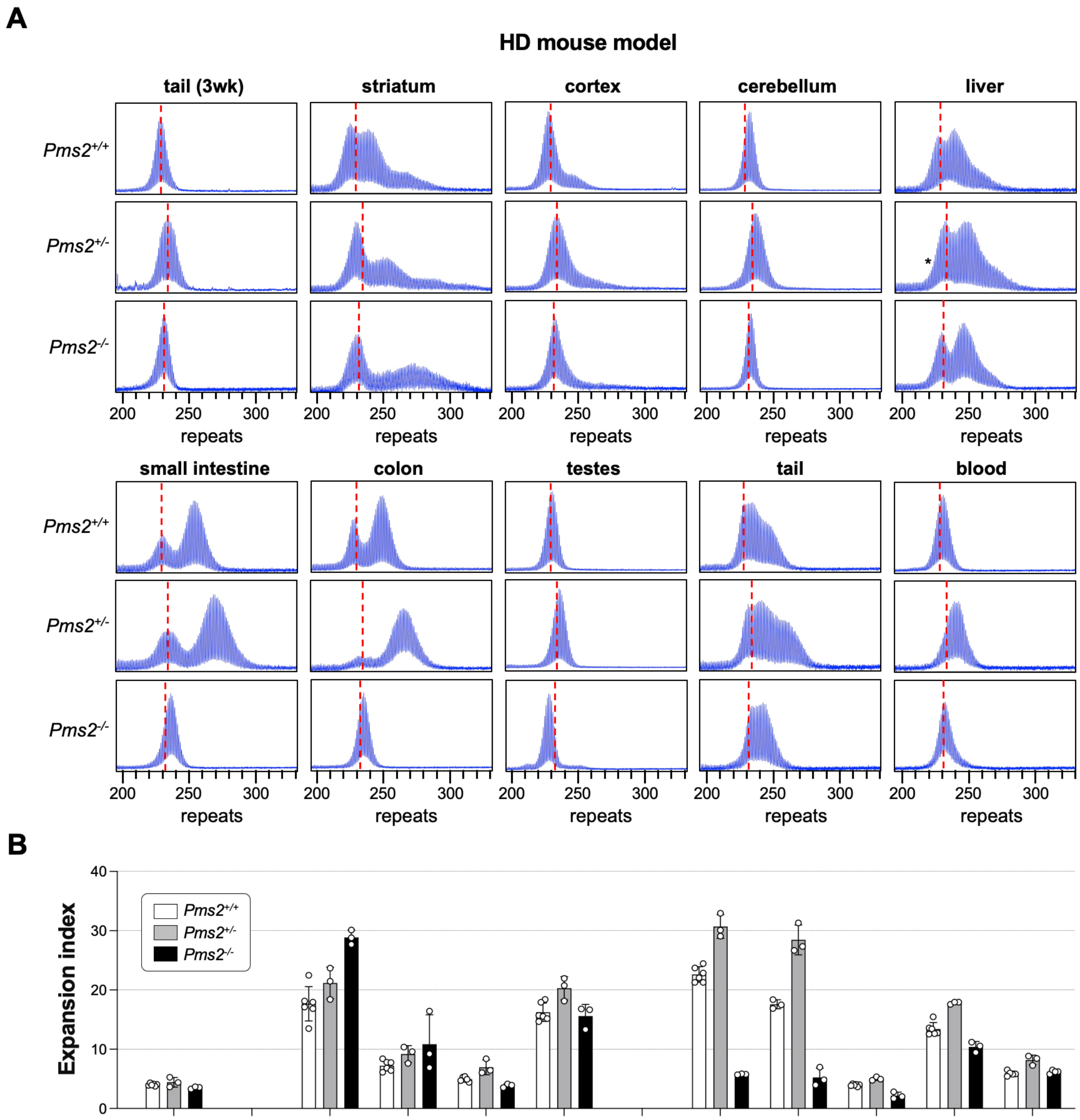


| | tail (3wk) | | striatum | cortex | cerebellum | liver | | small intestine | colon | testes | tail | blood |
|---|---|---|---|---|---|---|---|---|---|---|---|---|
| ***Pms2*$^{+/+}$ vs *Pms2*$^{+/-}$** | 0.9224 | | **0.0037** | 0.1562 | 0.1427 | **0.0007** | | **<0.0001** | **<0.0001** | 0.5669 | **0.0002** | 0.1018 |
| ***Pms2*$^{+/+}$ vs *Pms2*$^{-/-}$** | 0.8802 | | **<0.0001** | **0.0027** | 0.5733 | 0.8219 | | **<0.0001** | **<0.0001** | 0.2229 | **0.0156** | 0.9605 |
| ***Pms2*$^{+/-}$ vs *Pms2*$^{-/-}$** | 0.7351 | | **<0.0001** | 0.3719 | **0.0353** | **0.0007** | | **<0.0001** | **<0.0001** | 0.0572 | **<0.0001** | 0.1951 |

**Figure S3. The effect of *Pms2* deficiency on repeat expansion in different tissues from an HD mouse model.** (A) Representative repeat PCR profiles from tail DNA taken at 3 weeks (3 wk) and different organs of 4-month-old *Pms2*$^{+/+}$, *Pms2*$^{+/-}$ and *Pms2*$^{-/-}$ HD male mice with ~230 repeats. The dashed lines represent the sizes of the original inherited alleles as ascertained from the tail DNA taken at 3 weeks. (B) Comparison of the expansion index (EI) in the indicated organs of 4-month-old *Pms2*$^{+/+}$, *Pms2*$^{+/-}$ and *Pms2*$^{-/-}$ HD mice with an average of 234 repeats in the original allele. The colon data represent the average of 3 *Pms2*$^{+/+}$, 3 *Pms2*$^{+/-}$ and 3 *Pms2*$^{-/-}$ mice with 226-239 repeats. The data from other organs represents the average of 6 *Pms2*$^{+/+}$, 3 *Pms2*$^{+/-}$ and 3 *Pms2*$^{-/-}$ mice in the same repeat range. The error bars indicate the standard deviations of the mean. Each dot represents one animal. In each organ, the EIs for different genotypes were compared using a two-way ANOVA with correction for multiple testing as described in the Materials and Methods. The adjusted *p*-values are listed in the table below.

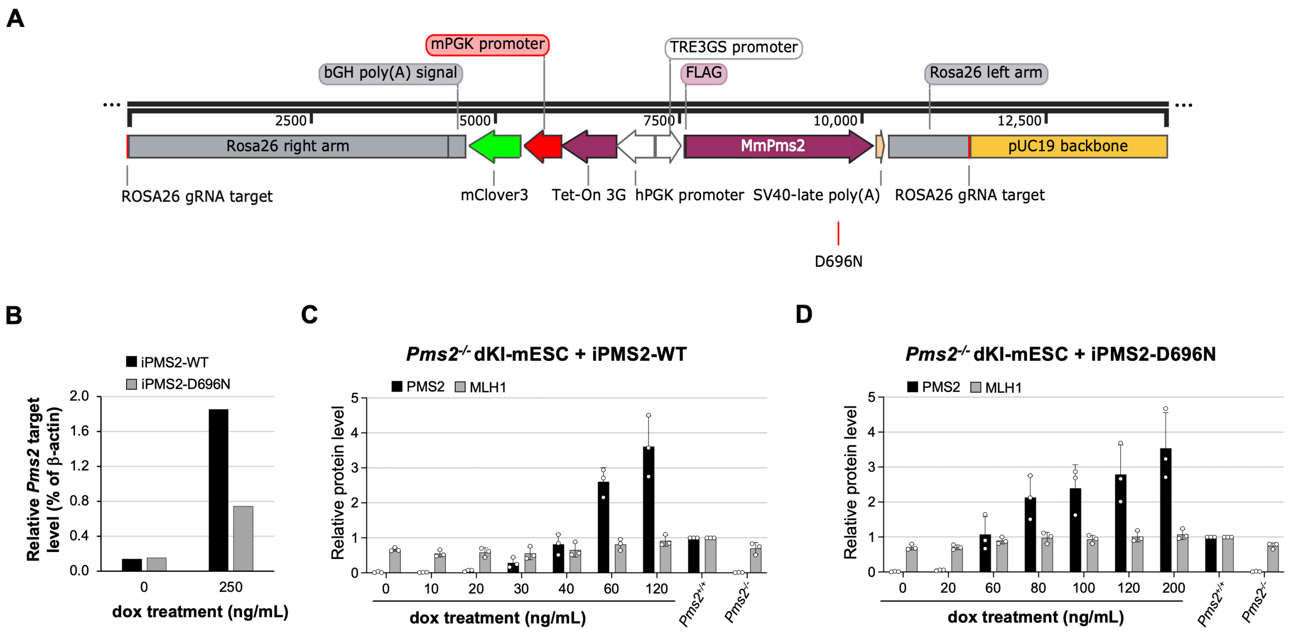


**Figure S4. Doxycycline (dox)-induced expression of FLAG-tagged *Pms2* constructs encoding either a WT (iPMS2-WT) or an endonuclease-deficient D696N mutant (iPMS2-D696N) form of PMS2.** (A) Linear representation of the constructs used to generate iPms2-WT and -D696N lines. PGK promoters drive constitutive expression of doxycycline-responsive Tet-On 3G protein and a mClover3 fluorescent marker. In the opposite orientation, the TRE3GS promoter drives doxycycline-inducible expression of *Pms2*. The D696N mutation is indicated in red below the *Pms2* cDNA sequence. (B) Quantitative PCR of *Pms2* targets, expressed as a percentage of β-actin, from mESC lines carrying both constructs and treated with the indicated concentrations of doxycycline. (C, D) The relative levels of PMS2 and MLH1 protein in mESC lines expressing the iPMS2-WT (C) and iPMS2-D696N (D) constructs. Quantitative analysis of Western blots of whole-cell lysates from mESCs treated with the indicated concentrations of doxycycline and from *Pms2*$^{+/+}$ and *Pms2*$^{-/-}$ control mESCs. Representative blots were shown in

Figure 4D and 5D. The uncropped blots showing the binding of antibodies to PMS2, MLH1, and β-actin are shown in Fig. S7. The band intensity was determined using Bio-Rad Image Lab v 6.1.0 build 7 software as described in the Materials and Methods. The PMS2 and MLH1 protein levels were normalized to β-actin, and the bar graph shows the fold change relative to the level in *Pms2*$^{+/+}$ cells. The data represents the average of 3 technical replicates. The error bars indicate the standard deviations of the mean. Each dot represents one replicate.

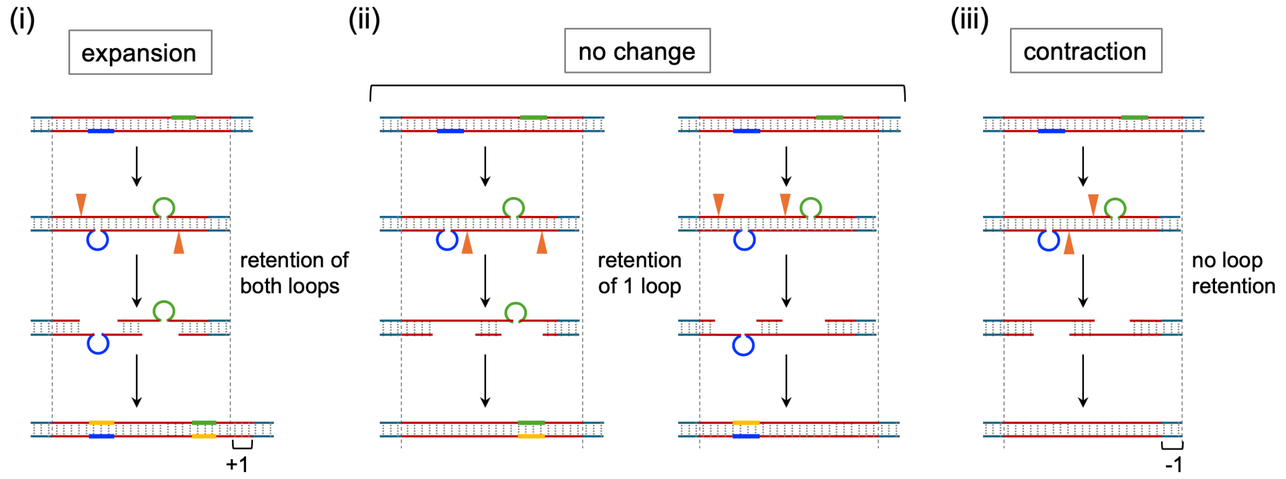


**Figure S5. Schematic representation of three possible outcomes for MutL-mediated processing of DNA substrates containing two loop-outs**. A double loop-out structure is suggested to form in the region of repeats when the DNA is transiently unpaired. Repeats are shown in red, and the blue and green lines indicate the repeats corresponding to the loop-outs. The yellow segments indicate the newly synthesized DNA produced using the loop-outs as a template. The orange triangles indicate MutL cut sites. (i) When both cuts occur on the strand opposite the loop-outs, as demonstrated for MutLγ-mediated cleavage *in vitro* (1), gap-filling will use the two looped-out regions as templates resulting in expansion. (ii) When both cuts occur on the same strand, excision or strand-displacement results in the removal of one loop-out and the incorporation of the other. After gap-filling the original allele will be restored (no change). (iii) In the case that both cuts occur on the same strand of the loop-outs, both loop-outs will be removed and result in contraction.

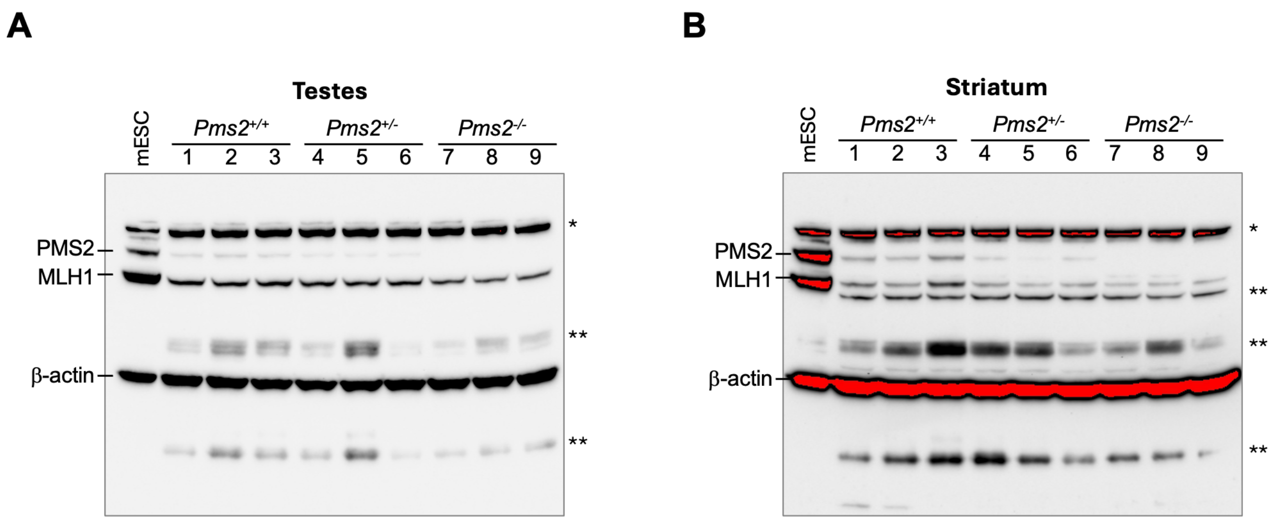


**Figure S6. The uncropped blots showing the binding of antibodies to PMS2, MLH1, and β-actin to representative mouse tissue.** Representative western blots showing the reaction of protein extracts from *Pms2*$^{+/+}$, *Pms2*$^{+/-}$, and *Pms2*$^{-/-}$ mouse testes (A) and striatum (B) with antibodies to PMS2, MLH1 and β-actin. Three different animals of each genotype are shown. Each lane of the testis and striatum samples contained 50 ng protein. A total of 20 ng of mESC samples were used as a positive control for these 3 antibodies. The normalization control, β-actin, is also shown on the same blot. A much lower exposure of this gel was used for the β-actin quantitation. The bands in red are heavily overexposed. The bands indicated by a single asterisk represent non-specific products resulting from the use of the MLH1 antibody. The bands indicated by two asterisks represent non- specific products resulting from the use of the PMS2 antibody.

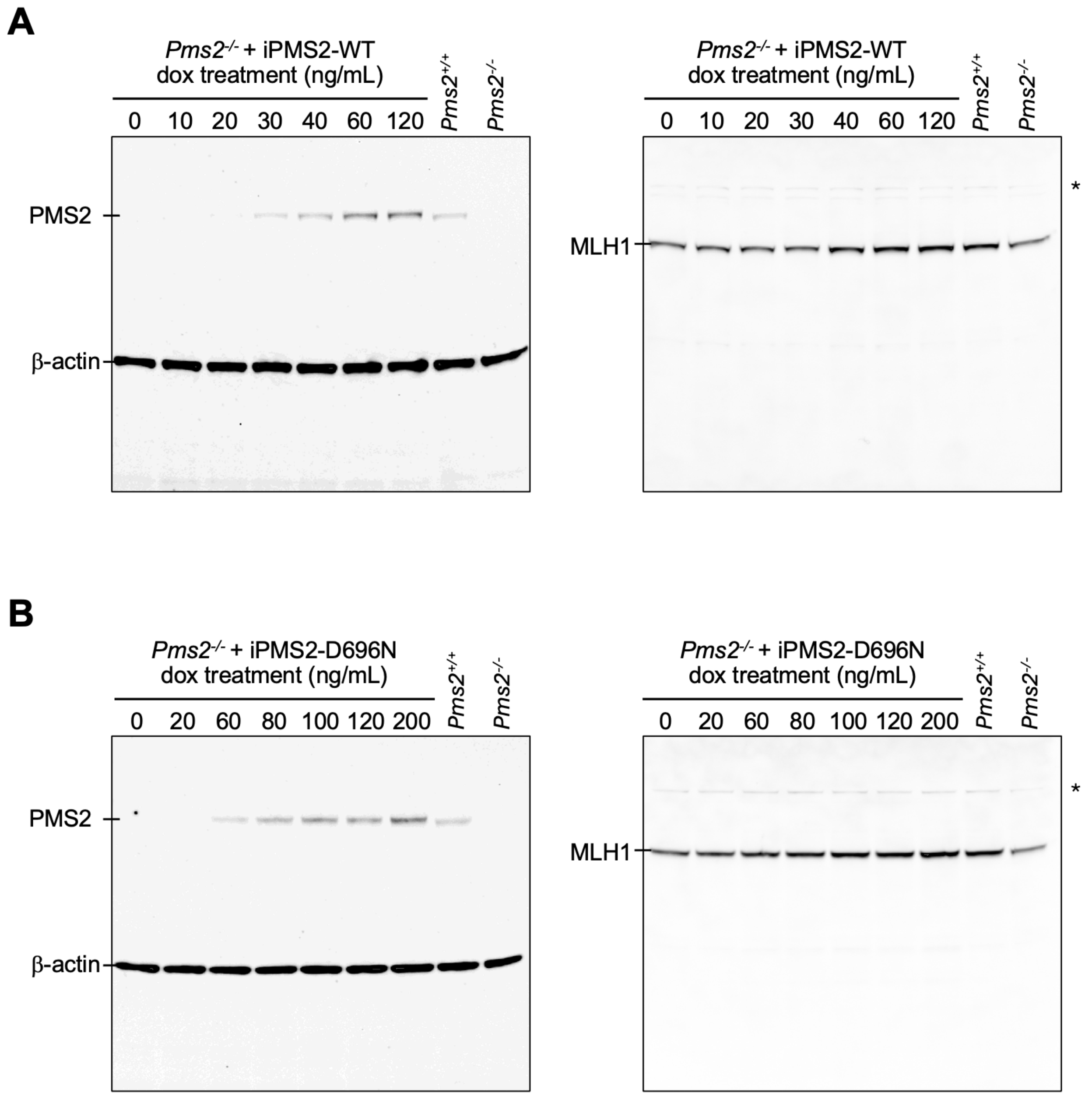


**Figure S7. Representative uncropped blots showing the binding of antibodies to PMS2, MLH1, and β-actin to extracts from m**ESCs expressing the iPMS2-WT (A) and iPMS2-D696N (B) constructs treated with different concentrations of DOX as indicate. Protein extracts from *Pms2*$^{+/+}$ and *Pms2*$^{-/-}$ mESCs were used as controls. The blots were probed sequentially with antibodies to PMS2 and MLH1 and the normalization control, β-actin. A much lower exposure of this gel was used for the β-actin quantitation. The bands indicated by a single asterisk represent non-specific products resulting from the use of the MLH1 antibody.

**References**


1. L. Y. Kadyrova, V. Gujar, V. Burdett, P. L. Modrich, F. A. Kadyrov, Human MutLgamma, the MLH1-MLH3 heterodimer, is an endonuclease that promotes DNA expansion. *Proc Natl Acad Sci U S A* **117**, 3535-3542 (2020).
